# Consequences of benzalkonium chloride tolerance for selection dynamics and *de novo* resistance evolution driven by antibiotics

**DOI:** 10.1101/2025.08.01.667674

**Authors:** Orestis Kanaris, Lydia-Yasmin Sobisch, Annett Gödt, Frank Schreiber, Niclas Nordholt

## Abstract

Biocides are used in large amounts in industrial, medical, and domestic settings. Benzalkonium chloride (BAC) is a commonly used biocide, for which previous research revealed that *Escherichia coli* can rapidly adapt to tolerate BAC disinfection, with consequences for antibiotic susceptibility. However, the consequences of BAC tolerance for selection dynamics and resistance evolution to antibiotics remain unknown. Here, we investigated in detail the effect of BAC-tolerance in *E. coli* on its response upon challenge with different classes of antibiotics. Competition assays showed that subinhibitory concentrations of ciprofloxacin - but not ampicillin, colistin and gentamicin - select for the BAC-tolerant strain over the BAC-sensitive ancestor at a minimal selective concentration of 0.0013-0.0022 µg/mL ciprofloxacin. In contrast, the BAC-sensitive ancestor was more likely to evolve resistance to ciprofloxacin, colistin and gentamicin than the BAC-tolerant strain when adapted to higher concentrations of antibiotics in a serial transfer laboratory evolution experiment. The observed difference in the evolvability of resistance to ciprofloxacin was partly explained by a negative epistatic interaction between the mutations conferring BAC-tolerance and a knockout mutation in the gene encoding for the outer membrane porin F. Taken together, these findings suggest that BAC-induced tolerance can be stabilized in environments containing low concentrations of ciprofloxacin, while it also constrains evolutionary pathways towards antibiotic resistance.

## Introduction

The widespread use of antiseptics and biocides has raised concerns regarding the emergence and spread of multidrug resistance^1^. However, there is limited understanding of the effect of antiseptics and biocides on the selection and development of antibiotic resistance^2^. Antiseptics and biocides are products that eliminate or inhibit harmful microorganisms. Both types of products may contain similar active substances, but they are used in different applications and are thus regulated by different laws in the EU^3,4^. Antiseptics are applied to living tissues, such as wounds or surgical sites, to prevent or treat infections^4^. In contrast, biocides are used in various environments, including industry, hospitals, agriculture, and household products, to control harmful organisms, such as disinfectants for inanimate surfaces and preservatives^1^. While antiseptics and biocides are in many cases based on active substances with a broad antimicrobial effect, active substances used in antibiotics as pharmaceuticals for the systemic treatment of humans or animals mostly rely on defined biochemical targets in bacteria.

Biocides used as disinfectants play a crucial role in medical and veterinary settings, especially since the worldwide rise of antibiotic resistance^5,6^. In the EU, 10660 tons of antibiotics were sold in 2017 for consumption by humans and for animals in the food-production industry^7^. This quantity is 40 times lower than the biocide sales in the EU, as estimated for 2009^7,8^. Due to the COVID-19 pandemic the use of biocides is increasing^9^. Antibiotic resistance has been related to biocide resistance^10^, and to exposure to subinhibitory concentrations of biocides^11^. Certain molecular mechanisms were shown to confer some levels of resistance to multiple drugs simultaneously, a phenomenon termed cross-resistance^12^. This highlights the need to investigate the consequences of the widespread use of biocides on resistance to antibiotics and biocides to identify risks of biocide use and to inform regulatory interventions.

Microorganisms are categorized to be resistant to a given antibiotic if therapy with the antibiotic will fail with high probability^13^. Such resistance is measured by determining the minimal inhibitory concentration (MIC), which is defined as the minimal antibiotic concentration inhibiting growth. Tolerance is a characteristic of a microorganism enabling it to survive treatment with an antimicrobial agent^14^. Tolerance is the main mode of microorganisms to evade the effects of disinfectants because the efficacy of disinfectants is based upon killing. Tolerance may be related to increased survival over many log-levels without a concomitant increase of the MIC^8^. Tolerant bacteria can slow or pause their growth in the presence of antimicrobials that require growth to harm the bacteria. While resistance and tolerance are often discussed together, they rely on distinct molecular mechanisms^14^, a point underscored by the observation that tolerance can increase without changes in MIC. For example, Zeng and colleagues^15^ demonstrated that repeated low-level biocide exposure selects for bacterial populations with complex survival strategies that may cross-protect against structurally and functionally different stressors, indicating potential links between biocide tolerance and multi-drug resistance. Moreover, it has been shown that antibiotic tolerance can facilitate the evolution of resistance to the same antibiotic^16^. Our previous work has shown that high levels of disinfectant tolerance evolve under periodic exposure^8,17^. These biocide-tolerant strains possess unique survival mechanisms that could have two consequences upon challenge with antibiotics; first, the decreased susceptibility may facilitate their selection in competition with sensitive strains and second, the differences in genetic make-up might impact their subsequent evolution of higher levels of resistance to biocides as well as antibiotics^18,19^. However, the response of biocide tolerant strains upon challenge with antibiotics has not been studied in detail.

Such above discussed consequences might be especially relevant in environments, in which biocides and antibiotics co-occur along with resistant or tolerant strains. Co-occurrence of antimicrobial substances like biocides and antibiotics happens in areas such as hospitals, animal husbandry, veterinary settings, and wastewater treatment^19,1,20^. It has been previously reported that bacteria that were in contact with the biocide benzalkonium chloride (BAC), developed a slightly reduced susceptibility to the antibiotics ampicillin (AMP) and ciprofloxacin (CIP) than before the exposure to BAC^8,21,22^. In addition, BAC-tolerant strains exhibited a higher growth rate as compared to their sensitive ancestral strain under subinhibitory concentrations of CIP^8^. Whether these slight fitness benefits alter the ability of the biocide-tolerant strains to be selected in mixed populations or if they facilitate the evolution of higher levels of antibiotic resistance it is not known.

Selection describes the potential of a specific genotype to dominate in numbers a population of different genotypes under specific conditions such as subinhibitory antibiotic concentrations ^23^. Increased fitness of resistant or tolerant strains over susceptible ones can lead to the selection, and subsequently to the dominance of the resistant or tolerant strains and the spread of antimicrobial resistance or tolerance in mixed populations. Under sub-MIC concentrations of antibiotics, the selection of resistant bacterial strains can be favored in direct competition over susceptible ones even at very low antibiotic concentrations (>230 times below the MIC)^23^. The lowest concentration of an antibiotic capable of selecting for the resistant genotype is known as the minimal selective concentration (MSC)^24^. Currently, it is unknown whether biocide tolerance provides a fitness advantage and thus selective benefits in the presence of antibiotics, and thereby is stabilized in the presence of antibiotics at environmentally relevant concentrations.

Biocide tolerant strains may not only be affected by antibiotics through differences in their competitive ability and thus selection. In addition, biocide tolerant strains may differ in their ability to evolve in the face of a challenge with high levels of antibiotics. The evolvability of resistance describes the ability of a microorganism to develop resistance to an antimicrobial agent and thereby maintain growth under lethal antimicrobial concentrations^25,26^. The evolvability of antibiotic resistance in biocide tolerant strains can be affected by changes in the mutation rate (i.e. the rate at which mutations occur) and epistatic interactions between genes or mutations that confer biocide tolerance with those genes or mutations that confer antibiotic resistance^27,28^. However, it is unknown whether tolerance to biocides affects the evolvability of antibiotic resistance. Determining the effects of evolved biocide tolerance on evolvability of antibiotic resistance is important for assessing the risks of biocide usage and subsequent emergence and fate of biocide tolerant strains, especially if those strains can cause infections that need to be treated with antibiotics.

In this study, we investigate the response of a BAC-tolerant *E. coli* strain upon challenge with the antibiotics CIP, AMP, colistin (COL), and gentamicin (GEN). Specifically, we focus on two distinct evolutionary processes: (i) the selection of the BAC-tolerant strain in competition with its susceptible ancestor in the presence of antibiotics and (ii) the *de novo* evolution of resistance to the four antibiotics in the BAC-tolerant strain as compared to its susceptible ancestor. To this end, we chose a BAC-tolerant strain that evolved in the laboratory in response to periodic disinfection cycles with lethal concentrations of BAC^8^ and compared it to its sensitive ancestor. Both strains were tagged with fluorescent proteins, and competition experiments were performed at different subinhibitory antibiotic concentrations. In addition, the BAC-tolerant and the ancestor strains were exposed to the four antibiotics in serial dilution adaptive laboratory evolution experiments with stepwise increasing antibiotic concentrations up to 2048 × the MIC to assess the effect of BAC-tolerance on the evolvability to antibiotic resistance. The experiments showed a clear effect of BAC tolerance on evolvability of CIP resistance. Whole-genome sequencing revealed that mutations in *ompF* were common in evolved ancestor strains but absent in BAC-tolerant populations, suggesting that epistatic interactions may underlie the observed differences in evolvability. This hypothesis was supported by targeted deletion of *ompF* in both the ancestral and BAC-tolerant backgrounds.

## Materials and methods

### Bacterial strains, media, culture conditions and chemicals

All strains used in this study are listed in (Table 1). *E. coli* MG1655 (thereafter referred as to the ancestor or wildtype, WT) and its laboratory evolved BAC-tolerant mutants (S1 to S6) used in this study, were routinely grown on M9 minimal medium (M9: 42 mM Na_2_HPO_4_, 22 mM KH_2_PO_4_, 8.5 mM NaCl, 11.3 mM (NH_4_)_2_SO_4_, 1 mM MgSO_4_, 0.1 mM CaCl_2_, 0.2 mM uracil, 1 µg/mL thiamine, trace elements (25 µM FeCl_3_, 4.95 µM ZnCl_2_, 2.1 µM CoCl_2_, 2 µM Na_2_MoO_4_, 1.7 µM CaCl_2_, 2.5 µM CuCl_2_, 2 µM H_3_BO_3_) and 20 mM glucose) at 37°C and 220 rpm. Where necessary, media were supplemented with ampicillin (AMP; Roth, K029.4), ciprofloxacin (CIP; Sigma, 17850-25G-F), gentamicin (GEN; Sigma, G1272-10mL), colistin sulfate (COL; Serva, 17420.02) or rifampicin (RIF; R3501-5G, Sigma-Aldrich). While introducing the fluorescent proteins to the *E. coli* strains, Luria-Bertani Lennox medium (LB medium) was used, which contained 10 g/L tryptone, 5 g/L yeast extract and 5 g/L NaCl. After the transformation of cells, super optimal broth (SOC medium) was used which contained 20 g/L tryptone, 5 g/L yeast extract, 0.5 g/L NaCl, 0.128 g/L KCL, 20 mM glucose and 40 mM MgSO_4_. The transposition of the transposon carrying the genes encoding the fluorescent proteins from the plasmids to the genomic DNA was induced by adding 0.1 % arabinose to LB medium. For serial dilutions, phosphate buffered saline (PBS) was used containing 10 mM (NH_4_)_2_SO_4_, 1.76 mM KH_2_PO_4_, 2.68 mM KCl and 137 mM NaCl. Plating and spotting were carried out on LB medium containing 1.5 % agar. Plates were incubated at 37 °C overnight unless stated otherwise.

**Table 1.**
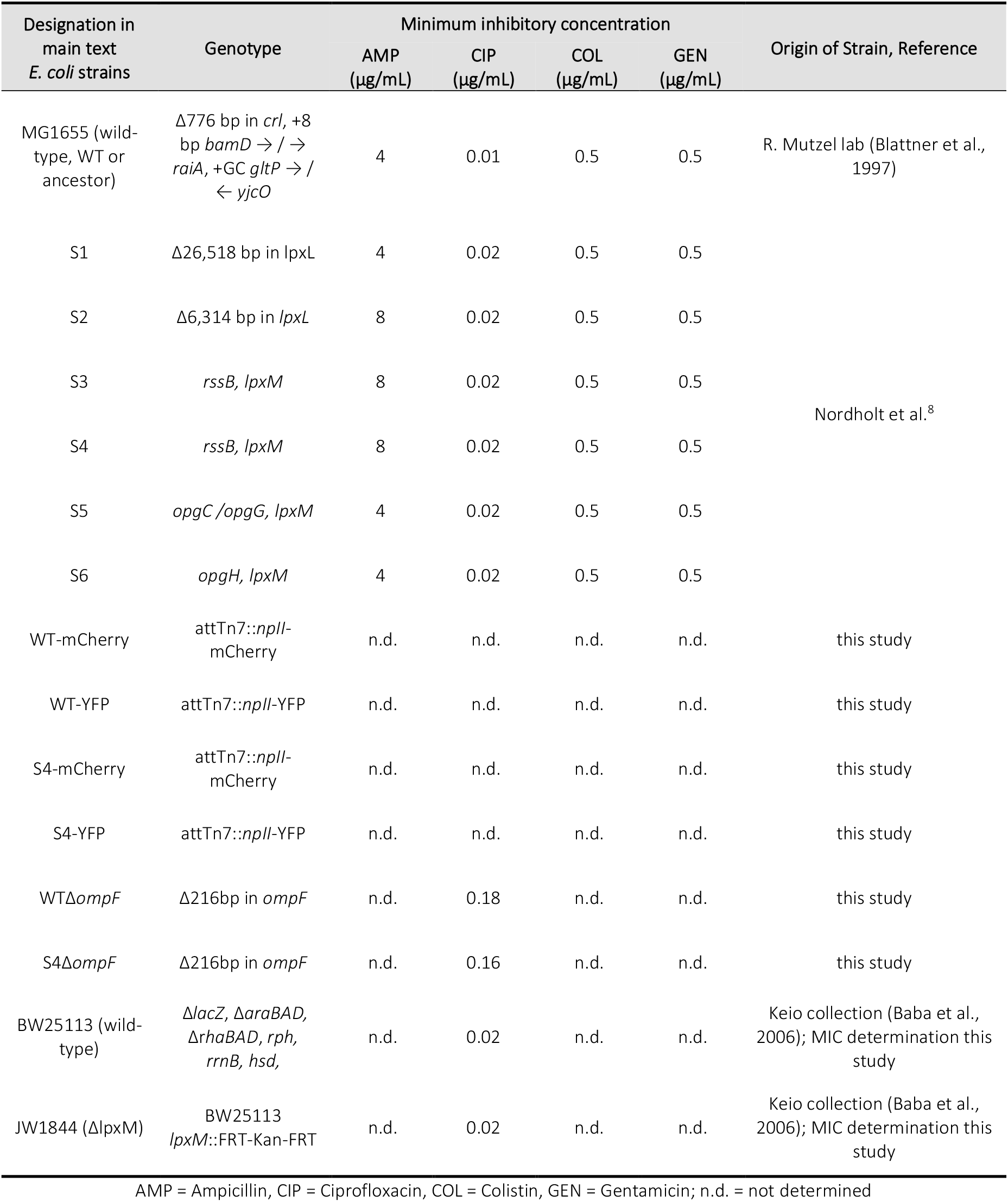
Strains used in this study including their genotype and their minimum inhibitory concentrations.

### Introducing fluorescent proteins into *E. coli* WT and biocide-tolerant *E. coli* S4

To prepare strains for the competition assays, plasmids pMRE-103 (mCherry), and pMRE-105 (YFP)^29^ were transformed into NEB 5-alpha chemically competent *E. coli* cells (NEB C2987I) following the manufacturer’s protocol with slight modifications. Briefly, 3 µL of plasmid was added to 50 µL of thawed cells, mixed gently, and incubated on ice for 30 minutes. After a 30-second heat shock at 42°C, cells were incubated in SOC medium at 30°C for 90 minutes. Cultures were then plated on LB agar medium with 100 µg/mL AMP and incubated overnight at 30°C. Plasmid isolation was performed using the Monarch Plasmid DNA Miniprep Kit according to the manufacturer’s instructions. The plasmid DNA was then quantified using a Nanodrop spectrophotometer (2000c, Peqlab). The plasmids were transformed into *E. coli* WT and BAC-tolerant *E. coli* strain S4 cells by electroporation. Cultures were grown to an OD600 of ∼0.3-0.6 in LB medium at 37°C with shaking. The cells were then washed with 10% glycerol. The washing step was repeated for four times and the cells were resuspended in ∼120 µL of 10% glycerol for storage or immediate use. Electroporation was performed using a BioRad GenePulser with a 2.5 kV pulse. After adding 3 µL of plasmid DNA, the cells were transformed and recovered in SOC medium at 30°C for 90 minutes. Transformed cells were plated on LB agar containing 100 µg/mL AMP and incubated overnight at 30°C.

For genome integration of genes encoding fluorescent proteins, the protocol from Remus-Emsermann et al. 2016^29^ was followed. Briefly, plasmid-containing bacteria were grown in LB with AMP and 0.1% arabinose to promote transposition. Colonies were screened for fluorescence with flow cytometry (Cytoflex S, Beckmann coulter) and confirmed by colony PCR using the attTn7 site-specific primers (5’GATGCTGGTGGCGAAGCTGT and 5’GATGACGGTTTGTCACATGGA)^30^. PCR was performed using OneTaq hot start master mix (M0484S, New England BioLabs) following the manufacturer’s protocol. The PCR cycle was set as following: initial 10 min step of 94°C, 30 cycles of 30 s 94°C, 30 s of 58°C and 105 s of 68°C. Next a final extension step of 5 min 68°C. The results were analyzed by 1% agarose gel electrophoresis (Mupid-exU).

### Generation of Δ*ompF* knockout strains

To analyse the role of *ompF* towards the evolvability of CIP resistance, knockout mutants in the WT and BAC-tolerant strain S4 were created by using the CRISPR-Cas9 system as described in Jiang et al. 2025^31^ by the company SYNBIO Technologies. The removal of selection markers introduced during cloning was confirmed by plating on plates with the appropriate antibiotics. The deletion was confirmed by amplifying the flanking regions of the deleted target site using PCR and sequencing. The sequence information is provided in Supplementary text 1. Aligning the amplified sequence with the *ompF* gene shows a 216 bp deletion in the *ompf* gene (base pairs 596-812).

### Determination of minimum inhibitory concentration (MIC) and dose-response curves for antibiotics

The MICs for CIP were determined using a modified version of the broth microdilution method^32^. Briefly, an exponentially growing culture was diluted to ∼10^7^ cfu/mL in a final volume of 200 μL M9 medium with increasing CIP concentrations. The following CIP concentration panels were used for different strains; JW1844 and BW25113: 0; 0.0025; 0.005; 0.01; 0.02; 0.04 µg/mL, S4 and WT: 0; 0.005; 0.01; 0.02; 0.04; 0.08 µg/mL, and S4Δ*ompF* and WTΔ*ompF:* 0; 0.0025; 0.005; 0.01; 0.02; 0.04; 0.08; 0.1; 0.12; 0.16; 0.18; 0.2 µg/mL. Strains were incubated in 96-well microplates (polypropylene (PP), Greiner Bio One) at 37°C for 24 h with shaking at 731 rpm in a BioTek Epoch 2 microplate reader. Optical densities were measured every five minutes and analyzed with the manufacturer’s software, Gen5 (version 3.09.). MIC values for the ancestor and strains S1 to S6 used in this study and additional antibiotics (CIP, AMP, COL, GEN) can be found in Nordholt et al.^8^. Dose-response curves were constructed by calculating the growth rates from the growth curves using the Python-based algorithm described in Swain et al. 2016^33^.

### Competition assay

Competitions between WT and biocide-tolerant strain S4 were conducted using WT-mCherry/S4-YFP and the reverse pair (WT-YFP/S4-mCherry) under four conditions per antibiotic in 96-well microtiter plates: M9 medium alone (control) and M9 medium with antibiotic (AMP: 0.5; 0.75; 0.85 µg/mL, CIP: 0.0025; 0.005; 0.01 µg/mL, COL: 0.2; 0.3; 0.4 µg/mL, GEN: 0.05; 0.1; 0.2 µg/mL). Nine overnight pre-cultures of each strain were grown in 200 µL M9 medium at 37°C with shaking at 220 rpm (Kuhner X Climo-shaker ISF1-X). Aliquots of 1 µL of both strains were then added into fresh 200 µL M9 medium with the appropriate antibiotic, thereby establishing a 1:200 diluted 1:1 mixed culture from which a sample was immediately taken to determine timepoint t=0. The cultures were incubated at 37°C under shaking at 220 rpm for 24 hours after which mixed cultures were then transferred into fresh medium with the appropriate concentration of antibiotics with a 1:100 dilution and grown again for 24 h. This cycle was repeated for three transfers in total. Samples were collected at the start of the experiment (t= 0, see above) at the end of each growth cycle (i.e. t= 24; 48; and 72 h), diluted appropriately in 0.9 % NaCl solution, and analyzed via flow cytometry (Cytoflex S, Beckmann Coulter) to quantify fluorescent populations. All media used in flow cytometry were filtered through a 0.1 or 0.2 µm filter. The samples were excited at 561 nm and measured at 585/42 nm to detect fluorescence of mCherry and excited at 488 nm and measured at 525/40 nm to detect fluorescence of YFP. The logarithm of the ratio of BAC-tolerant cells divided by the number of ancestor cells were plotted versus the number of generations of the competition. A linear regression was fitted to each replicate of the experiment to obtain the selection coefficients from the slope of the linear regression of these curves. Non-linear regressions were performed using a logistic model in R (drc package)^34^ to fit the selection coefficient over the antibiotic concentration data from different competitions treatments. Using these models, the MSC was determined as the antibiotic concentration at which the competitive advantage of the biocide-tolerant strain became apparent (i.e. the concentration at which the selection coefficient is estimated to be 0).

### Adaptive laboratory evolution experiment

We developed a design to investigate the effect of biocide tolerance on the probability of resistance evolution to antibiotics in a high-throughput evolution experiment. The fundamental concept of this design is the stepwise increase of antibiotic concentrations above the MIC in a large number of parallel cultures (96 to 348) and then to measure the fraction of cultures that are able to growth. Thereby, the experiment assesses the probability of each strain to evolve high-level resistance (i.e. their evolvability). The evolution experiments were conducted with the WT and the BAC-tolerant strains (S1 to S6) to evolve resistance against the antibiotic CIP. To investigate CIP resistance evolution in *lpxM* knockout mutant, we used *E. coli* JW1844 Δ*lpxM* (JW1844) and its parental strain *E. coli* BW25113 (BW25113) from the Keio collection^35^. In addition, separate evolution experiments with WT and BAC-tolerant strain S4 were conducted with the antibiotics AMP, COL, and GEN.

Cultures were grown in 96-deep-well plates (1.2 mL, Ratio lab), serially transferred every 48 hours at a 1:10 dilution in 700 µL final volume. The antibiotic concentration was initially set to 0.5 × MIC of the WT, increased to 4 × MIC and then increased 2 × to a maximum of 2048 × MIC. At each concentration step, 70 µL samples were taken to assess growth photometrically at 600 nm using a microplate reader (Epoch2, Biotek). Cultures were considered growing if their OD exceeded the average of blanks (media without cells) plus three times the standard deviation. In rare cases, growth in wells at late transfers were preceded by the absence of growth of the same lineage in earlier transfers. In this case, the lineage was retrospectively considered to have grown in the early transfers without being detectable. Considering the relatively low dilutions (1:10) at each transfer, and the final cell density of our *E. coli* WT in M9 of ∼10^9^ cells per 200 µL, statistically 9 transfers are necessary to exclude all the cells from a grown culture by dilution. It is possible that some cells acquire mutations that allow them to survive but only grow slowly and stay undetectable in the presence of the antibiotic. Subsequently such cells could acquire additional mutations to allow them to increase their growth rate, leading again to detectable growth. Stocks of cultures were prepared by adding glycerol to a final concentration of 15 % and storing them at -80°C.

### Sequencing of evolved populations

To investigate the molecular mechanisms behind CIP resistance evolution, DNA was extracted from 36 populations originating from the evolution experiments, including 18 WT populations and 18 populations of the BAC-tolerant strain S4. These populations have either adapted to the highest CIP concentration or not. From six populations per strain that adapted to the highest concentration, DNA was extracted at both the 64 × MIC and endpoint (2048 × MIC) concentrations. In addition, DNA was extracted only at 64 × MIC for six populations per strain that did not reach the endpoint. DNA extraction was performed using the peqGOLD Bacterial DNA Kit (VWR) from 300 µL of thawed stock. One population required regrowth under the original experimental conditions (2048 × MIC CIP) to yield sufficient DNA. Illumina whole-genome sequencing was conducted by Eurofins, and mutations were identified using the breseq pipeline^36^ in polymorphism mode to extract mutations with frequencies between 0.05 and 1. The genome NC_000913.3 (NCBI RefSeq accession) was used for the variant calling. The sequence data has been deposited in NCBI under Bioproject number PRJNA1282584.

### Mutation rate determination

Mutation rates of the BAC-tolerant strains were determined using the Luria-Delbrück fluctuation assay^37^. The strains were inoculated from frozen glycerol stocks into LB medium and then grown at 37°C with shaking at 220 rpm overnight. Overnight cultures were diluted to ∼4,000 cells/mL in fresh LB medium, and 54 independent cultures were inoculated with 200 µL of this dilution in 96-well plates. After growing the cultures overnight at 37°C, 220 rpm, the entire volume of 48 cultures was plated undiluted on LB agar plates supplemented with RIF (100 µg/mL) to determine the number of resistant mutants. Six cultures were diluted and plated on non-selective plates to estimate the total viable cell count per culture. After overnight incubation of the plates at 37°C for 24 h, the number of colonies was counted. This setup allowed for the estimation of mutations rates with the maximum likelihood method, using the rSalvador Package in R^38^. Statistical testing of differences between WT and BAC-tolerant strains S1 to S6 were performed with the Likelihood ratio test implemented in the rSalvador Package at p > 0.05. The mutation rate per base pair was calculated by dividing the maximum likelihood estimation by 79 which is the number of mutations known to confer resistance to RIF^39^.

### Statistics

Statistical analysis was carried out in R version 3.6.1. The data from the competition experiments of each antibiotic-concentration combination were tested for normality with the Shapiro-Wilk test and optically by visualizing the QQ-plots. Not all data passed the test for normality, therefore the non-parametric One-sample, two-sided Wilcoxon test was carried out to test if the data are significantly different from 0. The p-values were corrected for multiple comparisons with the method from Benjamini & Hochberg ^40^.

The data derived from the adaptive evolution experiments was used to calculate 95 % confidence intervals using the method from Clopper & Pearson^41^ for binomial data. Significance was assumed when the confidence intervals of the adaptive evolution experiments between the different strains were not overlapping.

Pearson’s Chi-squared test was used to test the difference in enrichment of mutations in genes in sequenced populations of the WT as compared to S4 at different stages of the adaptive evolution experiment. The p-values were corrected for multiple comparisons with the method from Benjamini & Hochberg^40^.

## Results & Discussion

### Subinhibitory concentrations of CIP select for BAC-tolerance

Competition experiments were conducted to understand the selection dynamics between the BAC-tolerant strain S4 and its ancestor *E. coli* strain (WT) under the influence of different antibiotics (AMP, CIP, COL, GEN). Both strains were fluorescently tagged (fluorescence proteins: YFP and mCherry) and analyzed in competition experiments using various subinhibitory concentrations of CIP, AMP, COL, and GEN. The experiments were carried out for both fluorescence combinations to exclude effects caused by the fluorescent proteins. Our results show that the WT dominates the population in the absence of antibiotics (Figure 1 and Supplementary Figure S1). The BAC-tolerant S4 strain exhibits a fitness advantage in the presence of increasing CIP concentrations, outcompeting the WT for both fluorescence combinations (Figure 1A and Supplementary Figure S1A). The MSC of CIP for the BAC-tolerant S4 strain was determined to range between 0.0013-0.0023 µg/mL, which corresponds to ∼4-fold lower than MIC (MIC_WT_=0.005-0.01 µg/mL). The competitive advantage of the BAC-tolerant strain S4 can be explained by its higher growth rate as compared to the WT in the presence of CIP^8^. In contrast, AMP, COL, and GEN did not select for the BAC-tolerant strain (Figure 1B-D). Even though GEN did not select for the tolerant strain consistently across both fluorescent combinations, the highest GEN concentration tested, led to very diverse outcomes between the replicates (Figure 1D and Supplementary Figure 1D). This indicates that subinhibitory concentrations of GEN close to the MIC can disturb the balance between different genotypes and lead to unpredictable population dynamics. The results of the competition experiments demonstrate that subinhibitory concentrations of certain antibiotics, such as CIP, can contribute to the selection of biocide-tolerant strains in direct competition with sensitive strains.

**Figure 1.**
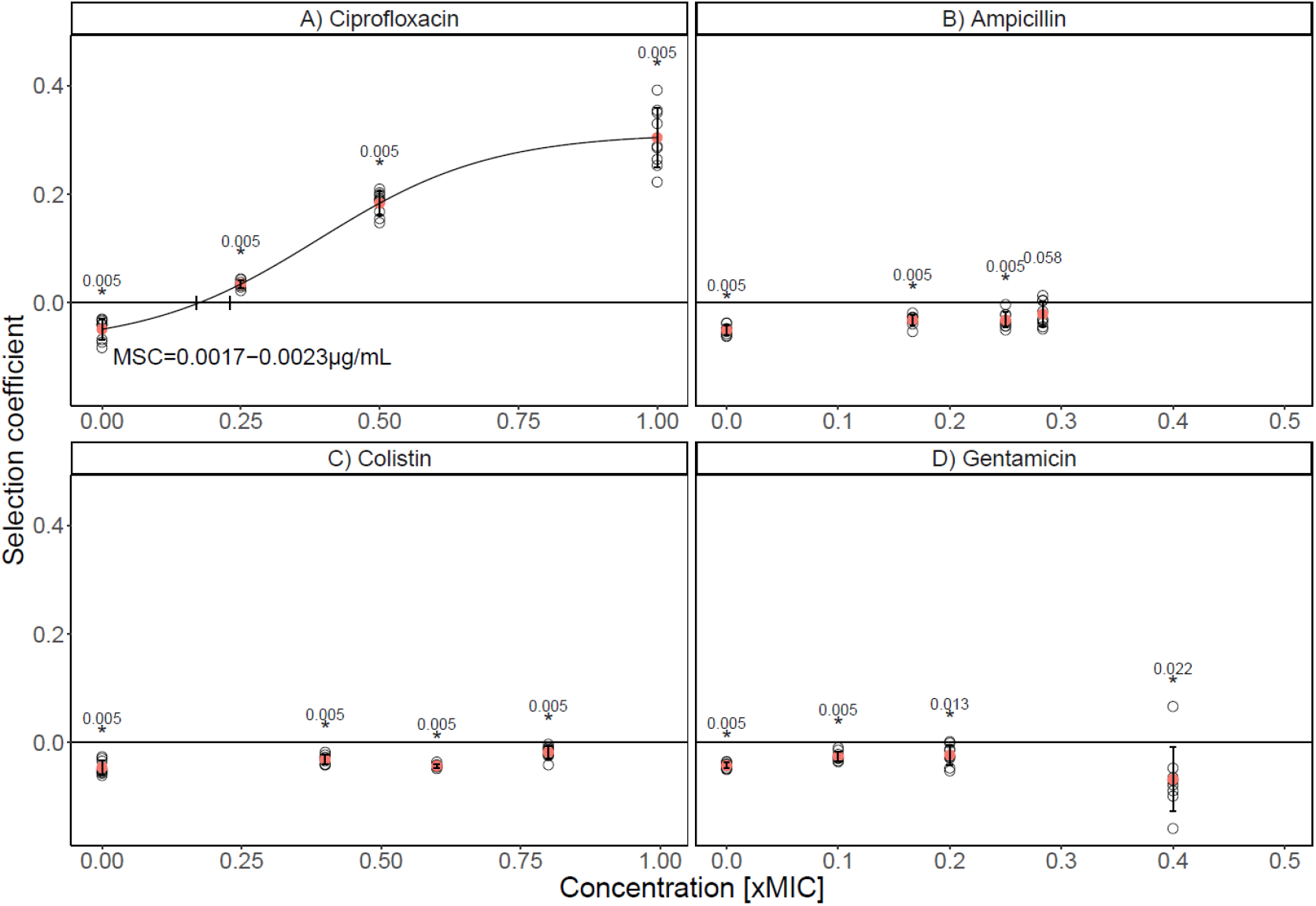
The benzalkonium chloride (BAC)-tolerant strain S4 has a selective advantage over the *E. coli* MG1655 wildtype/parental strain (WT) under ciprofloxacin (CIP) stress (A), but not in the presence of ampicillin (B), colistin (C) and gentamicin (D). Competitions between the WT and the BAC-tolerant strain S4 were performed under three different antibiotic concentrations (CIP: 0; 0.0025; 0.005; 0.01 µg/mL, AMP: 0; 0.5; 0.75; 0.85 µg/mL, COL: 0; 0.2; 0.3; 0.4 µg/mL, GEN: 0; 0.05; 0.1; 0.2 µg/mL). The panels display the selection coefficient of S4 against the WT as a function of the concentration represented as the fold-difference to the minimum inhibitory concentration (MIC) of the WT (CIP: 0.01 µg/mL; AMP: 4 µg/mL; COL: 0.5 µg/mL; GEN: 0.5 µg/mL). The selection coefficient was calculated from the change of S4 relative to the WT over generations of competition (as calculated from data shown in Supplementary Figure S2). The regression of the experimental results (black line) in A was fitted with a logistic function including all nine competitions per concentration (black, open circles). The intercept with the x-axis represents the minimum selection concentration (MSC). Positive values on the y-axis indicate selection for S4 and negative values selection for the WT. The red circles show the mean of the nine individual replicates (open circles). The error bars show the standard deviation. Competitions of WT-mCherry against S4-YFP are shown. Outcomes of competitions with reciprocal fluorescent tags are shown in Supplementary Figures S1 and S3. One-sample Wilcoxon test was performed against 0 and the p-values are shown above each concentration. Stars indicate significance (p<0.05) at n=9. The p-values were corrected for multiple comparisons with the method from Benjamini & Hochberg^40^.

The mutations in the BAC-tolerant strain S4 are not associated to high CIP resistance^8^. Nevertheless, in direct competition with the susceptible wild-type (WT), the BAC-tolerant strain exhibited a similar MSC to that of *E. coli* strains carrying clinically relevant CIP resistance mutations (e.g., *gyrA* D87N, Δa*crR*, and Δ*marR*). In previous studies, such resistance mutations were shown to have MSCs of approximately 0.1 of the MIC of the susceptible parental strain^23^. This suggests that the BAC-tolerant strain, despite not being resistant in terms of MIC, may possess a selective advantage under subinhibitory CIP concentrations against the WT. In Germany, CIP concentrations of similar range as the determined MSC (0.0013-0.022 µg/mL) were found in wastewater samples. Specifically, in hospital wastewaters and sewage sludge, maximum concentrations of ∼0.06 µg/mL and 0.6 µg/mL were reported, respectively, corresponding to approximately 30-fold and 300-fold higher than the MSC of the BAC-tolerant strain determined here. In addition, in pig and poultry slurry in Germany in 2018, a maximum of 0.1 µg/g CIP was recorded, i.e. approximately 50-fold higher than the determined MSC in this present study^42^. Considering that BAC is widely used as a disinfectant and cleaning agent in medical and animal husbandry, there is a risk that strains like the BAC-tolerant strain S4 could evolve in these environments. The concomitant presence of subinhibitory levels of CIP could alleviate the fitness costs of BAC tolerance and contribute to the selection of BAC-tolerant strains. As a possible consequence, disinfectant-tolerant bacteria could persist in these settings and constantly be released into the environment through the waste. Colonization with such bacteria could render BAC applied as disinfectant or antiseptic less effective.

### Tolerance to BAC limits evolvability of antibiotic resistance in a substance-dependent manner

Biocides and antibiotics may be applied in the same environment such as hospital or veterinary settings, and microorganisms may evolve tolerance to these biocides. Thus, it is important to understand how the evolution of biocide tolerance affects the evolvability of such strains in response to antibiotics. The evolvability of resistance to the antibiotics CIP, AMP, COL, and GEN for the BAC-tolerant strain S4 as compared to the WT was assessed by performing adaptive laboratory evolution experiments using 96 to 348 parallel serial dilutions every 48 h at increasing concentrations (from 0.5 x MIC to 2048 x MIC of the WT; Figure 2). This approach imposes a selective regime in which only resistant population proliferate. Regular testing of growth at each concentration step yields information on the probability of resistance evolution (i.e. the evolvability) by calculating the fraction of growing wells relative to all inoculated wells and plotting this against the increasing antibiotic concentration. The data shows that the WT has increased evolvability as compared to the BAC-tolerant strain S4 for CIP, COL, and GEN, while the evolvability to AMP was similar between both strains (Figure 2). Different BAC-tolerant strains were tested for their evolvability to CIP to investigate whether decreased evolvability to CIP was generally associated to BAC-tolerance. The data showed decreased evolvability of CIP resistance for all BAC-tolerant strains investigated (Supplementary Figure S4). However, the ability to evolve resistance against CIP between the different BAC-tolerant strains varied (decreased evolvability tendency: S4 > S3 > S5 > S6/S1 > S2).

**Figure 2.**
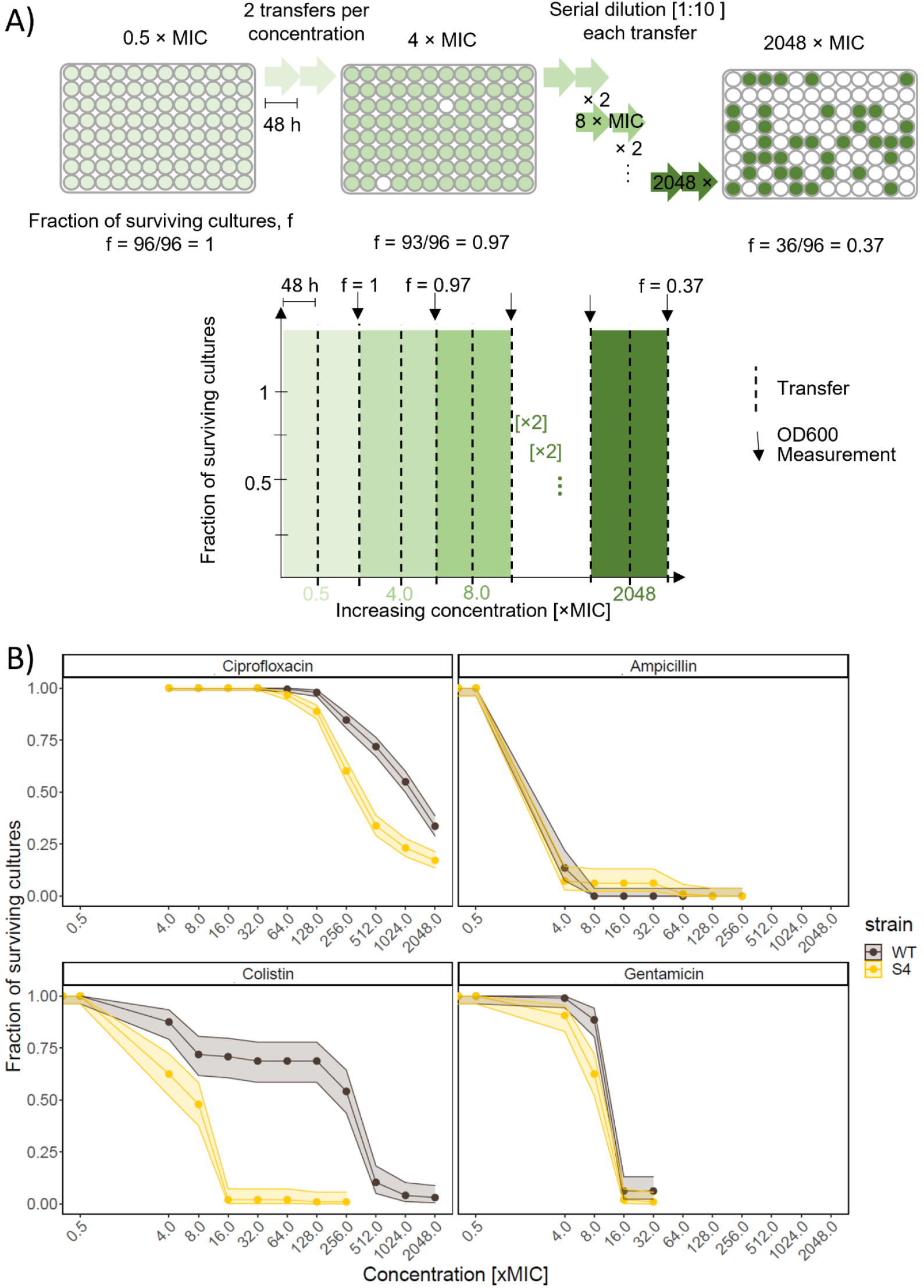
Benzalkonium chloride (BAC) sensitive *E. coli* MG1655 (wildtype, WT) shows increased evolvability against antibiotics as compared to its BAC-tolerant S4 strain. A) Scheme showcasing the design of the adaptive laboratory evolution experiment. The experiment was performed in 96-deep well plates. The cultures were initially incubated with a concentration of 0.5 × MIC, then the cultures were transferred to 4 × MIC, and thereafter the concentration was increased in 2-fold steps until 2048 × MIC. The cultures were transferred twice at each concentration and grown for 48 h after each transfer (dashed lines). OD600 was measured after the 2^nd^ transfer at any given concentration (arrows) and the fraction of growing (surviving) cultures f was determined. The cultures were diluted 1:10 at each transfer. B) The data shows the fraction of replicate evolving populations of each strain that had detectable growth at increasing concentration of antibiotics during the adaptive laboratory evolution experiment. Adaption to different antibiotics is shown for a concentration range of 0.5 to 2048 x MIC of the WT. Error bands show the 95 % confidence interval calculated using the method from Clopper & Pearson^41^ for binomial data. n=384 for ciprofloxacin and n=96 for the other antibiotics.

The difference between the strains in the response to CIP could be explained either by differences in mutation rates or by differences in accessible evolutionary paths between different genetic backgrounds of the BAC-tolerant strains as compared to the WT. Our data indicate that differences in mutations rates are not a likely explanation for the observed differences. Mutation rates µ were in a similar range between the BAC-tolerant strain S4 compared to the WT (µ_WT_ = 8.83∙10^−12^ mutations / bp / generation; µ_S4_ = 1,75∙10^−11^ mutations/bp/generation, p-value = 0.168 Likelihood ratio test; Supplementary Figure S5). Even the BAC-tolerant strains S3 and S6, which have significantly higher mutation rates than the WT (µ_S3_ = 2.72*10^−11^ mutations/bp/generation and p-value=0.014, Likelihood ratio test compared to the WT and µ_S6_ =2.53*10^−11^ mutations/bp/generation and p-value=0.016), were less likely to develop high CIP resistance. However, the mutation rates were measured in the absence of any antibiotics. It has been shown that subinhibitory concentrations of CIP increase the mutation rate of *E. coli*^27^. This could mean that the mutation rates of the strains changes during the experiment, which went undetected with our mutation rate assays.

The lack of detectable differences in mutation rates indicate that the differences in evolvability between strains can be explained due to differences in genetic background of the BAC-tolerant strains. All the BAC-tolerant strains have mutations in *lpxM* or *lpxL*, genes coding for acyl carrier proteins catalyzing the two last steps of Lipid A biosynthesis in the outer membrane ^8,43,44^. *LpxM* catalyzes the addition of a myristoyl group to lipid A, a key component of lipopolysaccharide (LPS) matrix, which is crucial for maintaining outer membrane stability, regulating membrane permeability, and establishing negative cell surface charge. Additionally, the S3 and S4 BAC-tolerant strains had a loss of function mutation in the regulator of the general stress response (*rssB*), S1, S5 and S6 a mutation in the synthesis of osmoregulated periplasmic glucans (*opGH*) and S2 had a deletion of genes related to mobility and chemotaxis^8^.

Since all BAC-tolerant strains S1-S6 show different nonsynonymous mutations in *lpxM* or *lpxL* genes, we hypothesized that these genes are involved in decreased evolvability to CIP and that their absence should affect evolvability. To test this hypothesis, we compared CIP evolvability between a *lpxM* knockout strain JW1844 from the KEIO collection and its ancestor *E. coli* BW25113. The results showed that deletion of the *lpxM* gene does not decrease evolvability to CIP (Supplementary Figure S4). The discrepancy in results may be explained by the fact that the BAC-tolerant strains carry point mutations in *lpxM* or *lpxL*, which could alter protein function without fully inactivating it. In contrast, strain JW1844 carries a complete knockout of *lpxM*, potentially leading to more drastic phenotypic effects. Alternatively, the observed result may be specific to the BW25113 background.

### Deletions in the *ompF* gene are significantly enriched in WT populations as compared to BAC-tolerant populations during adaptation to CIP

We sequenced the whole genomes of 36 populations from the evolution experiment to obtain a deeper understanding on the role of the genetic background of BAC-tolerant strains affecting evolutionary paths towards CIP resistance. To this end, we sequenced populations from the adaptive laboratory evolution experiments originating from WT and S4. We selected six populations originating from the WT as well as S4 that successfully adapted to the highest concentration of the experiment and six that went extinct before reaching the highest CIP concentration of the experiment. Of the populations that adapted to the highest concentration, we sequenced the DNA at two concentration-steps, 64 × MIC and endpoint of 2048 × MIC. Of the populations that went extinct before the endpoint, we sequenced the DNA only from the 64 × MIC concentration step. Across all populations, a total of 70 genes at 132 positions were mutated with a frequency in the population of 20 % or higher. All sequenced populations had at least one mutation in the *gyrA* gene with 100 % frequency, except for one population that had a mutation in *gyrB* instead of *gyrA* (Figure 3). The genes *gyrA* and *gyrB* encode for the DNA gyrase subunits A and B, respectively, which is the primary target of fluoroquinolone antibiotics in *E. coli*^45^. Statistical testing identified one gene (*ompF*) that was mutated with higher prevalence in populations originating from the ancestor strain as compared to populations originating from S4 (Pearson’s chi-squared test, p=0.0273, n=6, adjusted for multiple comparisons with the Benjamini & Hochberg, 1995 method^40^). The gene *ompF* encodes for the outer membrane porin F. Mutations in *ompF* were significantly enriched in WT-populations compared to BAC-tolerant-populations both at 64 x MIC and 2048 x MIC.

**Figure 3.**
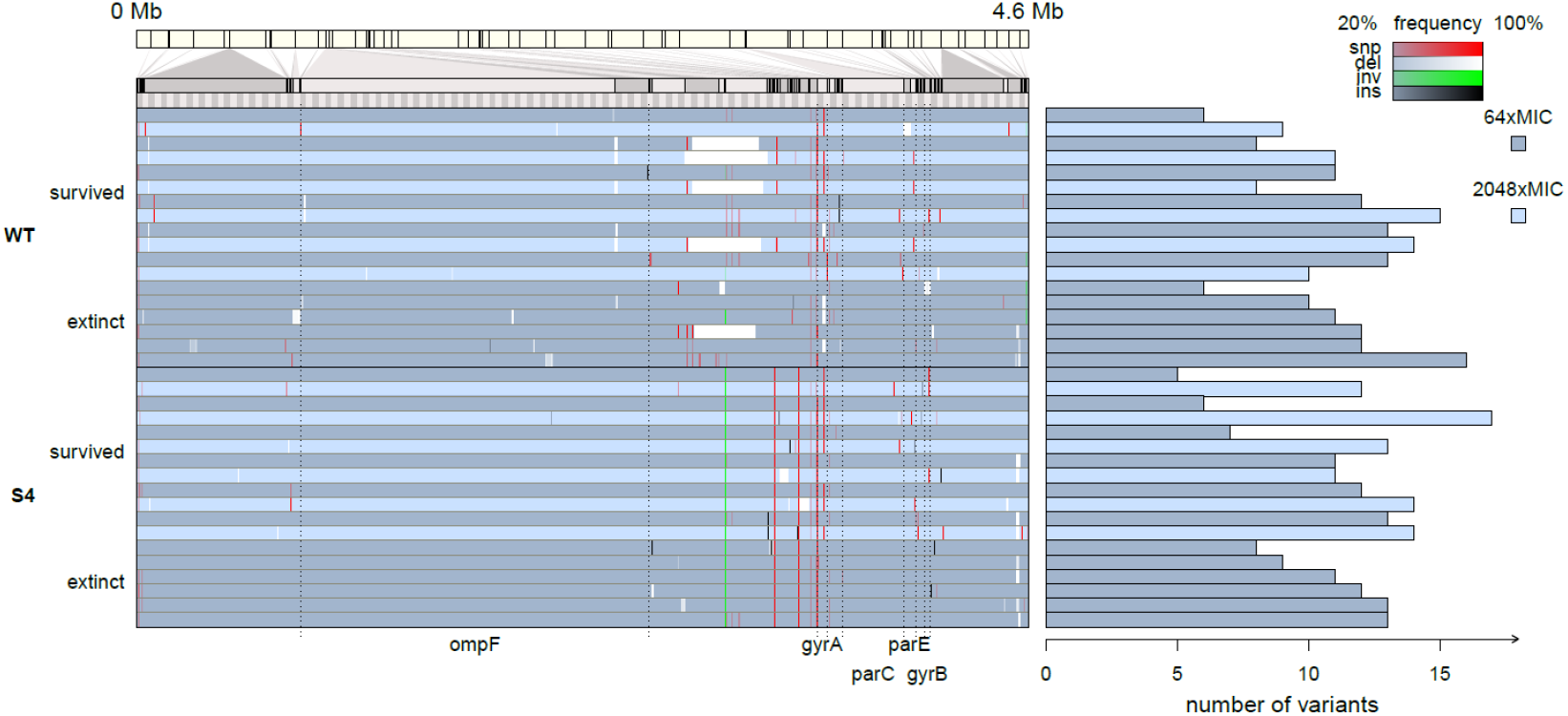
The *ompF* gene is preferentially mutated in the WT strain and not in the BAC-tolerant strain upon adaptive laboratory evolution to ciprofloxacin (CIP). Whole genome sequence analysis of *E. coli* MG1655 (WT) and its BAC-tolerant derivative S4 after experimental evolution in the presence of CIP. The bar at the top represents the genome of *E. coli* from 0 MB to 4.6 Mb. Genomic loci with mutations are magnified in the panel below. Each of the 36 blue bars represent one replicate *E. coli* population from the evolution experiment. Dark blue bars show populations from the 64 × MIC concentration step of the experiment. The populations that adapted (‘survived’) to the highest concentration (2048 × MIC) of the experiment are represented by light blue bars. A dark blue bar (64 × MIC) followed by a light blue bar (2048 × MIC) represents sequences from a single evolved population at the two different concentrations. For both, the WT and S4, 6 populations that survived the entire evolution experiment (i.e. they adapted to 2048 × MIC) are shown. In addition, the 6 dark blue bars at the bottom of each strains’ segment (WT and S4) represent genomes of evolved populations at 64 × MIC that went extinct at concentration above 64 x MIC and below 2048 × MIC (‘extinct’). Red lines represent single nucleotide polymorphisms (snp), white lines represent deletions (del), green lines represent inversions (inv) and black lines represent insertions (ins). The intensity of the line shows the frequency of the specific mutation in the population. Names of key genes are indicated below the diagram. The number of variants in each population is shown on the right side of the figure.

The whole genome sequencing data indicates that *ompF* mutations shape adaptation to CIP and that differences in *ompF*-based adaptation leads to differences in evolvability at high CIP concentrations. All the mutations in *ompF* were either deletions or insertions of length not multiple of three, causing a frameshift and consequently most likely rendering the gene inactive. The absence of OmpF was shown to contribute partially to resistance against another fluoroquinolone, norfloxacin, and was associated with a lower accumulation of norfloxacin inside of *E. coli* cells^46^. Although the MIC increase to CIP conferred by a *ompF* knockout is relatively low^47^, it is known that low resistance mutations can have positive epistatic effects with the major fluoroquinolone resistance mutations^48^. The WT populations that survived up to 2048 × MIC displayed four different mutations in the *gyrA* gene at 64 × MIC (D87G, Δ3 bp nucleotides: 247-249, S83L and G81D), while the BAC-tolerant strain had only two mutations in *gyrA* (D87G and S83L). This difference was not statistically significant in our experiment, but it is an indication that the occurrence of the knockout in *ompF* provides access to evolutionary pathways that enable specific mutations in the *gyrA* gene that are not accessible without the *ompF* loss-of-function. It should be noted that the high number of mutated genes compared to the number of sequenced populations affects the statistical power of the analysis. Thus, it is possible that other genes are differently mutated in the two strains but were not detected by our analysis.

### *ompF* loss-of-function has negative epistatic interactions with the BAC-tolerance mutations

We hypothesized that the loss-of-function mutations in *ompF* are not selected in the S4 strain because of negative epistatic interactions with the existing BAC tolerance mutations present in the S4 strain. To this end, we obtained knockouts of the *ompF* gene in the WT and the S4 BAC-tolerant backgrounds. We then measured dose-response curves for CIP of both strains. The results show that the absence of the *ompF* gene increased the MIC (∼8-fold) of the WT, as reported previously ^47^, but only slightly increased the MIC of the BAC-tolerant strain (∼2-fold). Strikingly, fitness costs of the *ompF* deletion were substantially higher in the BAC-tolerant strain S4 as compared to the wild-type (61 % vs 17 % fitness loss, respectively, Figure 4). These results indicate a negative epistatic interaction between the absence of *ompF* and the mutations related to BAC-tolerance. Epistasis is widespread in nature, and it can be an important driver of the direction of evolution^49^. Epistatic effects can be both positive and negative. Negative epistasis was detected in the context of antibiotic resistance acquisition^28^. Even multiple individually beneficial mutations can have an overall negative epistatic effect and reduce the fitness of a strain^50,51^. Our data shows that biocide tolerance mutations can exhibit negative epistasis with antibiotic resistance mutations.

**Figure 4.**
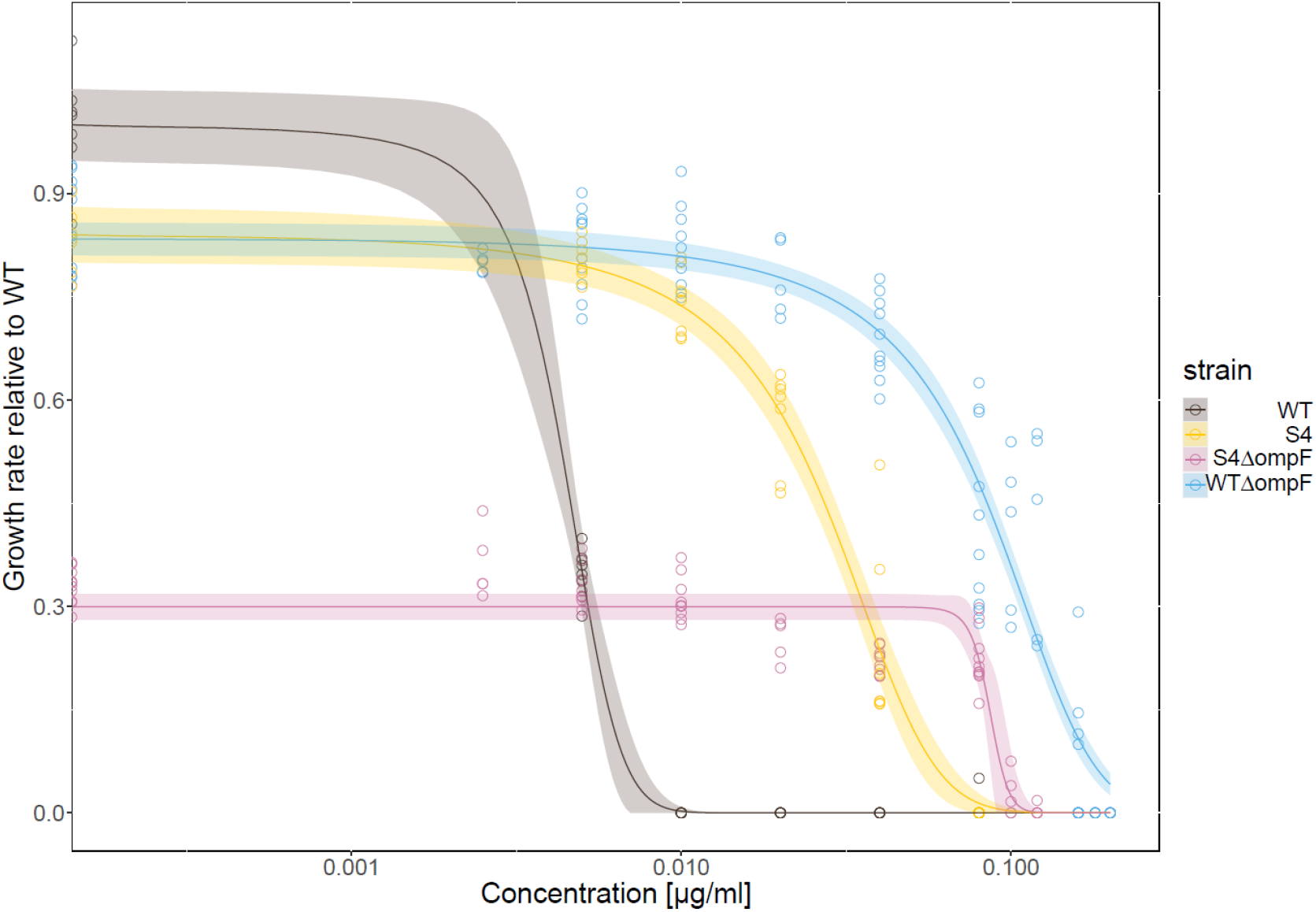
Dose-response curves to ciprofloxacin (CIP) of *E. coli* MG1655 (wildtype, WT), the BAC-tolerant S4 strain and their *ompF* knockouts. The y-axis shows the growth rate of the strains relative to the growth rate of the WT without CIP. The x-axis shows the CIP concentration on a logarithmic scale. The WT is represented in brown color, S4 in yellow, S4Δ*ompF* in pink and WTΔ*ompF* in blue. The lines show a logistic fit of the data, fitting 3 parameters: upper limit, the half-maximal inhibitory concentration (EC50) and slope. The lower limit was fixed to 0. The shaded area represents the 95 % confidence interval according to the fit. The MIC values of the used strains as measured here are WT: 0.005-0.01 µg/mL. S4: 0.04-0.08 µg/mL, WTΔ*ompF*: 0.16-0.18 µg/mL, S4Δ*ompF:* 0.12-0.16 µg/mL.

A reduced expression of *ompF* was observed in *E. coli* clinical isolates and from animals^47,52,53^, suggesting that it is a common pathway to high CIP resistance and not an artefact of our experimental design. In clinical isolates of *Klebsiella spp.*, a reduced expression of major porins was also observed^54^. *E. coli ompF* knockout mutants also showed slightly increased MICs to β-lactam antibiotics, tetracycline, chloramphenicol, and norfloxacin due to lower accumulation of the antibiotics in the cells^46,55^.

We note that solving the molecular mechanism of the negative epistasis was beyond the scope of our work, but linking our results to the literature allows to formulate hypotheses. All BAC-tolerant strains used for the evolution experiments have altered LPS. Negative interactions between the altered LPS and the absence of OmpF could explain the observed difference in antibiotic resistance evolvability between the BAC-tolerant strains and the BAC-sensitive WT. A large part of the surface of the outer membrane of *E. coli* is covered by β-barrel protein arrays next to patches of LPS ^56^. OmpF is a β-barrel protein and one of the major outer membrane porins (OMP) in *E. coli* ^56,57^. The absence of OmpF proteins could allow larger areas of the outer membrane to be covered by LPS and act as a barrier preventing the uptake of CIP^56^. The BAC-tolerant strain, having a mutation in LPS biosynthesis, possesses a different LPS composition. This altered composition in combination with the absence of OmpF could impair the function of the membrane, as the interactions between LPS and OMP are crucial for the integrity, stability, and assembly of the outer membrane^58^.

The second mutation that is present in the BAC-tolerant strain S4 is a loss-of-function in *rssB*. RssB regulates the degradation of RpoS, the master regulator of the general stress response ^59,60^. Non-functioning RssB leads to accumulation of RpoS in the cell, which results in the sustained activation of the general stress response ^60^. RpoS regulates the expression of a plethora of genes in *E. coli* associated with adaptation to different stresses, protein processing and cell physiology^60,61^. Genes that are differently expressed as regulated by RpoS could also have negative epistatic interactions with the *ompF* loss-of-function mutation and could therefore be responsible for the observed differences between the BAC-tolerant strain S4 and the WT.

Taken together, our data provide evidence for a negative epistasis between mutations conferring BAC tolerance and mutations in *ompF*. The reduced evolvability connected with this negative epistasis suggest that there is a common evolutionary pathway to high CIP resistance through the loss-of-function of *ompF*, which is blocked for the BAC-tolerant strain, because of the negative epistasis, and thus the BAC-tolerant strain is less likely to evolve high CIP resistance.

## Conclusions

Biocides and antibiotics such as BAC, AMP, GEN, COL, and CIP are used jointly or present in environments like hospitals, animal husbandry, and wastewater. Thus, BAC tolerance can evolve in these settings and the evolved BAC-tolerant bacteria can be exposed to antibiotics. Once BAC tolerance emerges, subinhibitory concentrations of CIP can select for BAC-tolerant *E. coli* strains over susceptible strains in those settings, and lead to the release of such strains into the environment. Contaminations with such strains could compromise the efficacy of BAC used for disinfection. Moreover, such BAC-tolerant *E. coli* strains may cause infections, which need to be treated with antibiotics. Our data shows that these BAC-tolerant strains are less likely to adapt to antibiotic concentrations that are relevant for such antibiotic treatments. One mechanistic explanation for this is through impaired access to a loss-of-function of *ompF* during evolution of CIP resistance. The deletion of *ompF* is not accessible in the BAC-tolerant strain because of negative epistasis between the BAC tolerance mutations and the loss-of-function of *ompF*. In turn, the acquisition of the *ompF* deletion is a part of the evolutionary pathway to high CIP resistance. Taken together, our study highlights that the evolution of biocide tolerance will have intricate effects on the selection dynamics of such strains in the presence of antibiotics and their potential for *de novo* evolution of antibiotic resistance.

## Data availability

DNA sequences generated here can be found in NCBI under Bioproject number PRJNA1282584. All other data generated during this study are included in this article and its supplementary information files.

## Acknowledgements

We thank Jan Kreft and Jens Rolff for discussions, and Mitja Remus-Emsermann for sharing plasmids. O.K. acknowledges funding by the Deutsche Bundesstiftung Umwelt (DBU; #20022/012-34/2). This work was supported by the Federal Institute for Materials Research and Testing within the funding scheme ‘Menschen Ideen – Typ 1’ (#MIT1-17-13).

## Credit authorship contributions

**O.K.:** Conceptualization, Methodology, Validation, Formal analysis, Investigation, Data Curation, Writing -Original Draft, Writing – Review & Editing, Visualization, Funding acquisition; **L.Y.S.** Validation, Formal analysis, Investigation, Data Curation, Writing - Original Draft, Writing – Review & Editing, Visualization, Supervision; **A.G.:** Formal analysis, Investigation; **F.S.:** Conceptualization, Methodology, Resources, Writing - Original Draft, Writing - Review & Editing, Supervision, Project administration, Funding acquisition; **N.N.:** Conceptualization, Methodology, Software, Writing - Review & Editing, Supervision, Funding acquisition.

