## Supplementary information for "Consequences of benzalkonium chloride tolerance for selection dynamics and *de novo* resistance evolution driven by antibiotics"

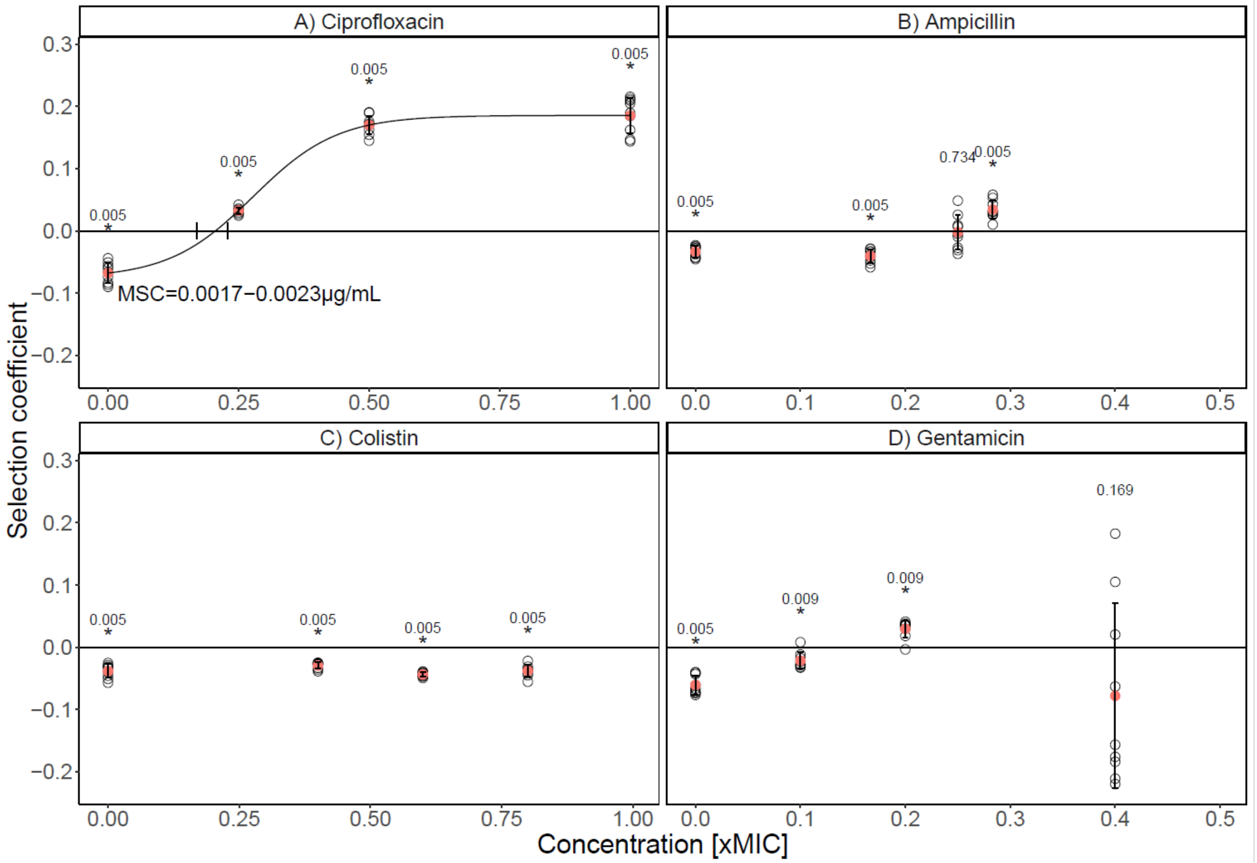


**Supplementary Figure S1. The benzalkonium chloride (BAC)-tolerant strain S4 has a selective advantage over the *E. coli* MG1655 wildtype/parental strain (WT) under ciprofloxacin stress (A), but not in the presence of ampicillin (B), colistin (C) and gentamicin (D).** Competitions between the WT and the BAC-tolerant strain S4 were performed under three different antibiotic concentrations (CIP: 0; 0.0025; 0.005; 0.01 µg/mL, AMP: 0; 0.5; 0.75; 0.85 µg/mL, COL: 0; 0.2; 0.3; 0.4 µg/mL, GEN: 0; 0.05; 0.1; 0.2 µg/mL). The panels display the selection coefficient of S4 against the WT as a function of the concentration represented as the fold difference to the minimum inhibitory concentration (MIC) of the WT (CIP: 0.01 µg/mL; AMP: 4 µg/mL; COL: 0.5 µg/mL; GEN: 0.5 µg/mL). The selection coefficient was calculated from the change of S4 relative to the WT over generations of competition (as calculated from data shown in Supplementary Figure S3). The regression of the experimental results (black line) in A was fitted with a logistic function including all nine competitions per concentration (black, open circles). The intercept with the x-axis represents the minimum selection concentration (MSC). Positive values on the y-axis indicate selection for S4 and negative values selection for the WT. The red circles show the mean of the nine individual replicates (open circles). The error bars show the standard deviations Competitions of WT-YFP against S4-mCherry are shown. Outcomes of competitions with reciprocal fluorescent tags are shown in Figure 1. One-sample Wilcoxon test was performed against 0 and the p-values are shown above each concentration. Stars indicate significance ($p<0.05$) at $n=9$. The p-values were corrected for multiple comparisons with the method from Benjamini & Hochberg[40].

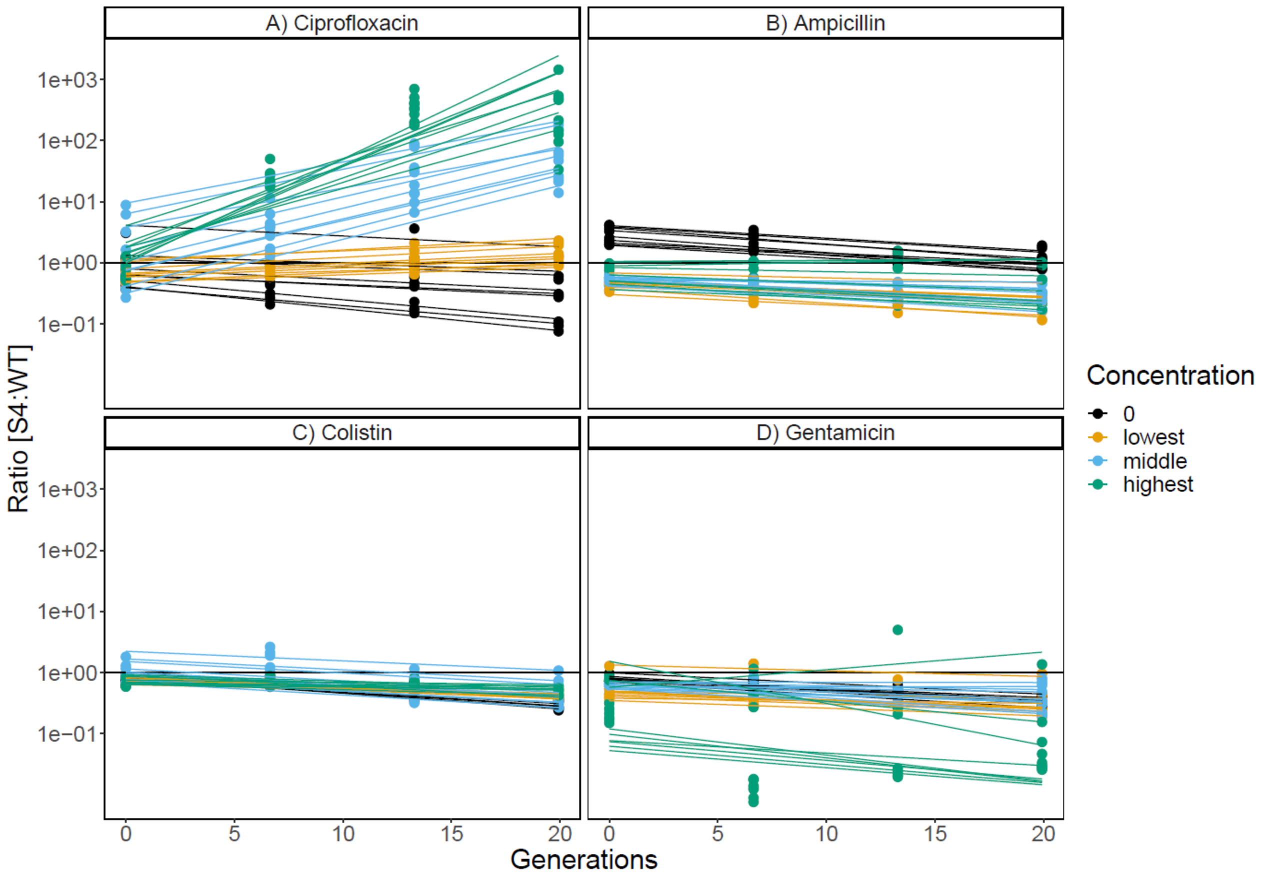


**Supplementary Figure S2.** Population dynamics of the competition experiment in the presence of four different antibiotics between the wildtype (WT) strain expressing mCherry and the BAC-tolerant strain S4 expressing YFP. Each panel shows the ratio of S4 divided by the WT plotted against the generations of the competition experiment. The Y axis is logarithmic. Lines represent linear regressions fitted for each replicate of the experiment (n=9). The colors represent the different antibiotic concentrations. In black, the results in the absence of antibiotics are shown, in yellow, the lowest antibiotic concentration, in blue, the middle concentration used and in green the highest. The antibiotic concentrations used in the competition experiments are CIP: 0.0025; 0.005; 0.01 µg/mL, AMP: 0.5; 0.75; 0.85 µg/mL, COL: 0.2; 0.3; 0.4 µg/mL, GEN: 0.05; 0.1; 0.2 µg/mL. The slopes of the linear regressions are the selection coefficients shown in Figure 1.

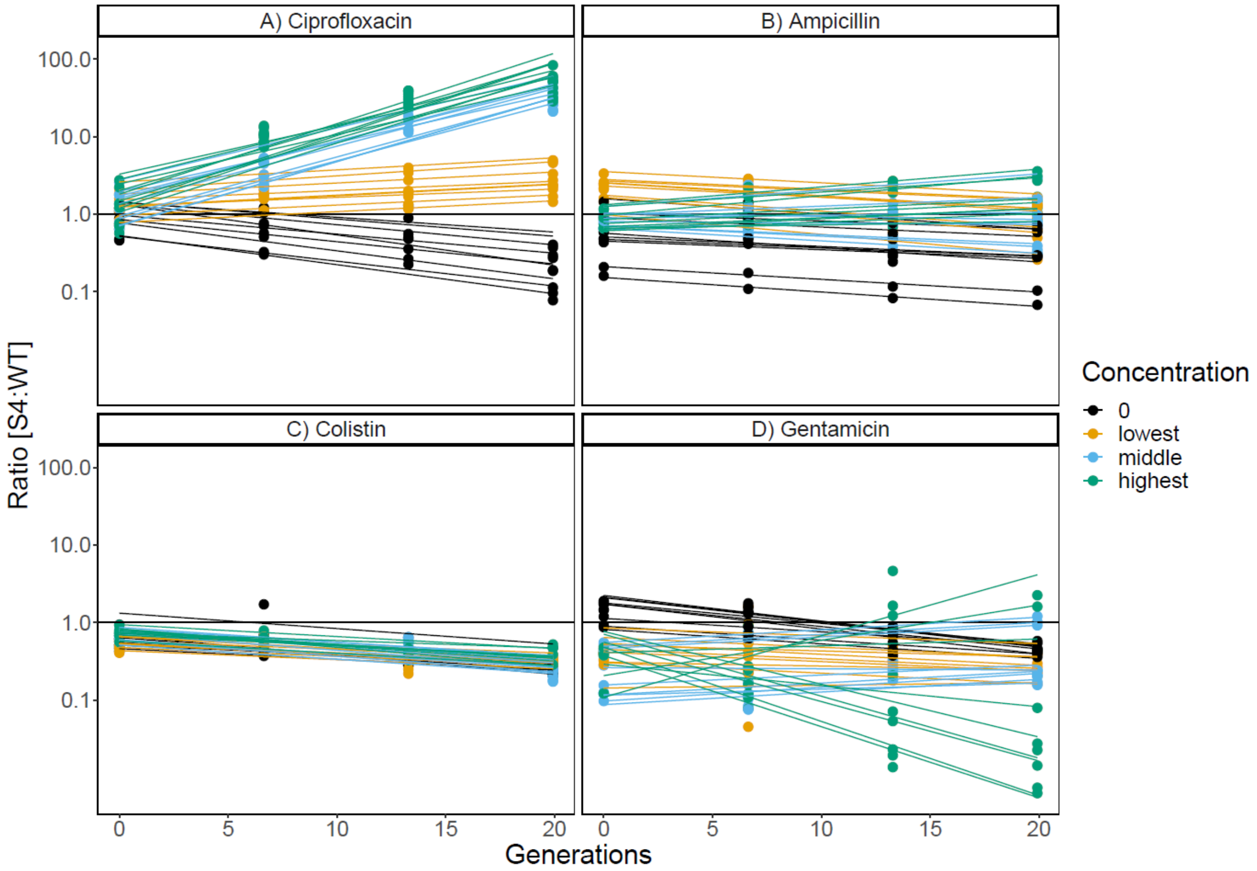


**Supplementary Figure S3.** Population dynamics of the competition experiment in the presence of four different antibiotics between the wildtype (WT) strain expressing YFP and the BAC-tolerant strain S4 expressing mCherry. Each panel shows the ratio of S4 divided by the WT plotted against the generations of the competition experiment. The Y axis is logarithmic. Lines represent linear regressions fitted for each replicate of the experiment (n=9). The colors represent the different antibiotic concentrations. In black, the results in the absence of antibiotics are shown, in yellow, the lowest antibiotic concentration, in blue, the middle concentration used and in green the highest. The antibiotic concentrations used in the competition experiments are CIP: 0.0025; 0.005; 0.01 µg/mL, AMP: 0.5; 0.75; 0.85 µg/mL, COL: 0.2; 0.3; 0.4 µg/mL, GEN: 0.05; 0.1; 0.2 µg/mL. The slopes of the linear regressions are the selection coefficients shown in Supplementary Figure S1.

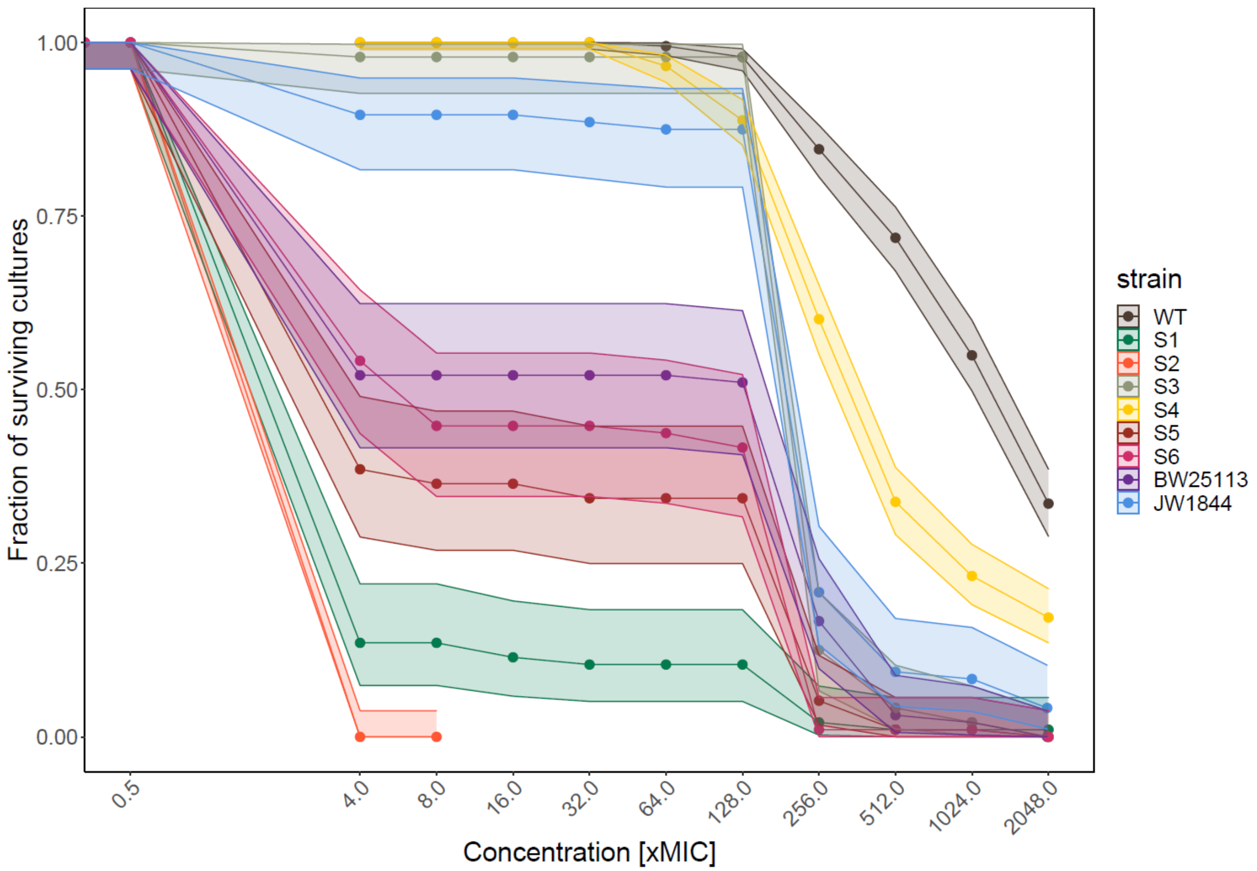


**Supplementary Figure S4. Benzalkonium chloride (BAC) sensitive *E. coli* MG1655 (wildtype, WT) shows increased evolvability against ciprofloxacin as compared to different laboratory-evolved BAC-tolerant strains.** The data shows the fraction of replicate evolving populations of each strain that had detectable growth at increasing concentration of antibiotics in a serial transfer adaptive laboratory evolution experiment. Adaption to different antibiotics is shown for a concentration range of 0.5 to 2048 x MIC of the WT. The strains S1-S6 are BAC-tolerant, JW1844 has a knockout in lpxM and BW25113 is its parental strain. Error bands show the 95 % confidence interval calculated using the method from Clopper & Pearson[41] for binomial data. n=384 for WT and S4 and n=96 for all other strains.

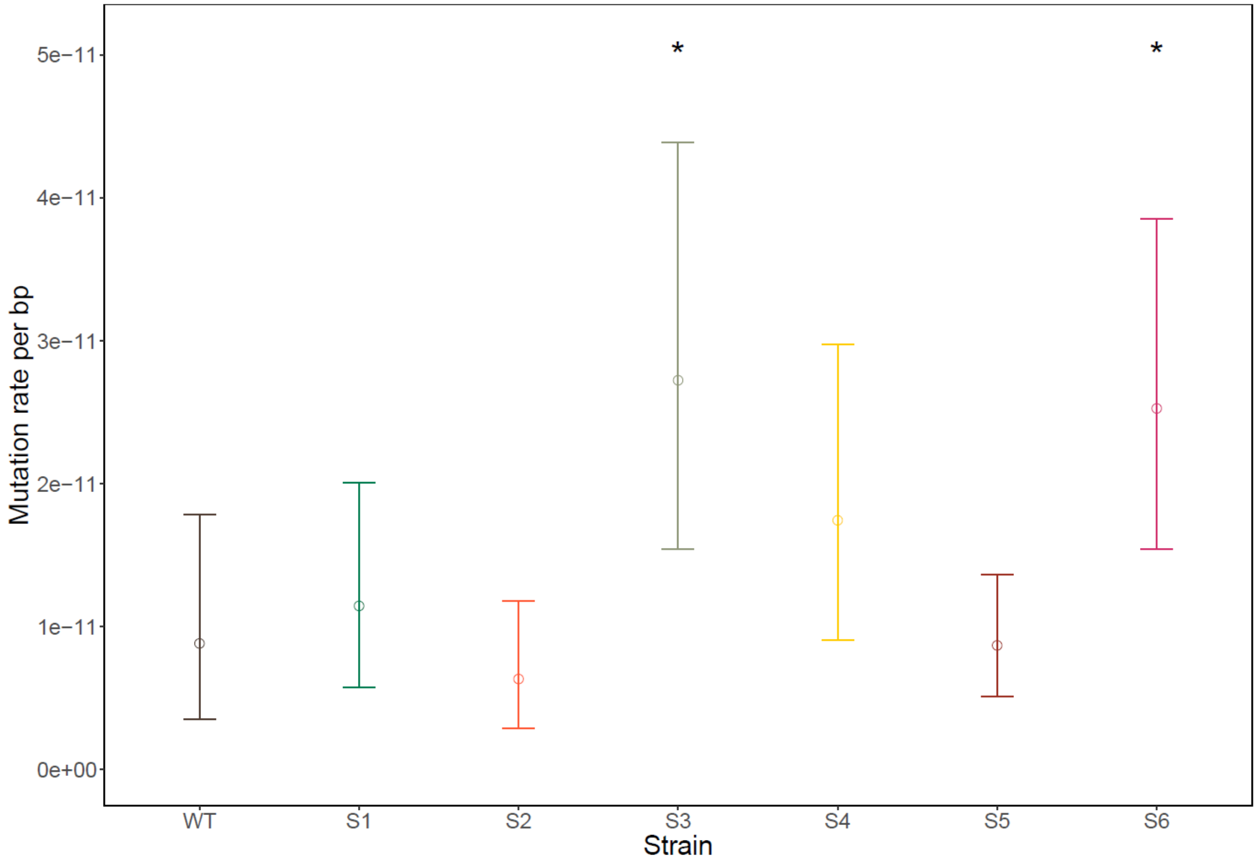


**Supplementary Figure S5.** Mutation rates of the WT and the BAC-tolerant strains. The data show the per base pair rates for each strain estimated by maximum likelihood method. The error bars represent the 95 % confidence intervals. The stars designate the strains with a statistically significant difference in mutation rate compared to the WT ($p < 0.05$) as tested with Likelihood ratio test (S3 $p = 0.014$ and S6 $p = 0.016$).

**Supplementary text 1**

The sequence flanking the deletion in the *ompF* gene in the strain WTΔ*ompF* is:

____ACCGATTCCTTTCTTCCGTCTTTCGCTTTAGATTTGGTGTAAGCGATGGACGGACGCAGACCGAAATCGAACTG
GTATTGCGCAACTAACAGAACGTCTTGCGTTTTGTTGGCGAAGTTAACCGCCAACACCGTCGCCGTTAGAACGGCGT
GCAGTGTCACGCTCGTTTTTACCCAGGTACTGAACAGCGAAGTTCAGGCCATCAACCAGACCAAAGAAGTTGGAGTT
ACGATAGGTAGCAACGCCGCCAACACGACCAACGAAGGGGTCATCGC____

The highlighted base indicates the region of the deletion of a 216 bp segment from the original *ompF* gene.

The sequence flanking the deletion in the *ompF* gene in the strain S4ΔompF is:

____CCAGTATAACTTCCCGATTCCTCATTACGTCTCCCGCTATCAGGATTTGGTGTAAGCGATGGACGGACGCAGACC
GAAATCGAACTGGTATTGCGCAACTAACAGAACGTCTTGCGTTTTGTTGGCGAAGTTAACCGCCAACACCGTCGCCG
TTAGAACGGCGTGCAGTGTCACGCTCGTTTTTACCCAGGTACTGAACAGCGAAGTTCAGGCCATCAACCAGACCAAA
GAAGTTGGAGTTACGATAGGTAGCAACGCCGCCAACACGACCAACGAGGAGATCATCGC___

The highlighted base indicates the region of the deletion of a 216 bp segment from the original *ompF* gene.
